# Pheniqs 2.0: accurate, high performance Bayesian decoding and confidence estimation for combinatorial barcode indexing

**DOI:** 10.1101/2021.03.11.434956

**Authors:** Lior Galanti, Dennis Shasha, Kristin C. Gunsalus

## Abstract

**Background:** Systems biology increasingly relies on deep sequencing with combinatorial index tags to associate biological sequences with their sample, cell, or molecule of origin. Accurate data interpretation depends on the ability to classify sequences based on correct decoding of these combinatorial barcodes. The probability of correct decoding is influenced by both sequence quality and the number and arrangement of barcodes. The rising complexity of experimental designs calls for a probability model that accounts for both sequencing errors and random noise, generalizes to multiple combinatorial tags, and can handle any barcoding scheme. The needs for reproducibility and community benchmark standards demand a peer-reviewed tool that preserves decoding quality scores and provides tunable control over classification confidence that balances precision and recall. Moreover, continuous improvements in sequencing throughput require a fast, parallelized and scalable implementation.

**Results:** We developed a flexible, robustly engineered software that performs probabilistic decoding and supports arbitrarily complex barcoding designs. Pheniqs computes the full posterior decoding error probability of observed barcodes by consulting basecalling quality scores and prior distributions, and reports sequences and confidence scores in Sequence Alignment/Map (SAM) fields. The product of posteriors for multiple independent barcodes provides an overall confidence score for each read. Pheniqs achieves greater accuracy than minimum edit distance or simple maximum likelihood estimation, and it scales linearly with core count to enable the classification of >11 billion reads in 1h15m using <50 megabytes of memory. Pheniqs has been in production use for seven years in our genomics core facility.

**Conclusions:** We introduce a computationally efficient software that implements both probabilistic and minimum distance decoders and show that decoding barcodes using posterior probabilities is more accurate than available methods. Pheniqs allows fine-tuning of decoding sensitivity using intuitive confidence thresholds and is extensible with alternative decoders and new error models. Any arbitrary arrangement of barcodes is easily configured, enabling computation of combinatorial confidence scores for any barcoding strategy. An optimized multithreaded implementation assures that Pheniqs is faster and scales better with complex barcode sets than existing tools. Support for POSIX streams and multiple sequencing formats enables easy integration with automated analysis pipelines.

## Background

High-throughput next-gen bulk sequencing with multiplexed sample barcodes is now standard practice to increase throughput and reduce sequencing costs, and single-cell applications are rapidly proliferating and evolving [1]. The advent of barcodes to tag individual cells and molecules from which sequence reads originate both enhances resolution and helps control for quantification biases. Yet indeterminate reads that are discarded, and uneven barcode distributions that differ from expectation, are common and decrease both accuracy and sensitivity.

The fast pace of innovation in sequencing technologies and experimental designs continuously presents new challenges to the classification of sequence reads. Increasingly complex barcoding schemes are being devised to accommodate novel experimental designs, and combinatorial cellular indexing protocols involving several successive rounds of barcoding expand the potential space exponentially [2] [3]. Just over the horizon, multimodal profiling – the simultaneous measurement of gene expression, protein abundance, chromatin state, spatial transcriptomics, and/or CRISPR-based readouts (e.g. lineage tracing, genetic screens) – is poised to become a new industry standard [1]. Third-generation platforms overcome some of the limitations of short-read sequencing by providing very long reads, but current long-read sequencers operate with very high error rates of 15-40%. To compensate, these are often complemented with high-fidelity short-read sequences or self-corrected using many-fold coverage [4]. Multiplexing with barcoding is now emerging as a way to increase throughput and lower costs for these platforms as well.

Several tools are currently available for decoding and classifying barcodes, an essential task for demultiplexing pooled bulk sequence libraries and for assigning reads to individual cells in single-cell workflows. However, existing tools for decoding index tags do not provide flexible support for the rising complexity of novel barcoding schemes. As a result, numerous custom solutions to handle different barcoding schemes are being implemented downstream of standard demultiplexing software, often geared to a specific application and implemented using convenient scripting languages such as R or Python. Various integrated workflows focusing on scRNA-seq analysis now bundle identification of cellular barcodes and molecular barcodes (UMIs) with transcript quantification and other downstream analyses to identify and study different cell types, including CellRanger [5], Seurat [6], Salmon-Alevin [7], and BUStools [8]. Such workflows could benefit from preprocessing by an efficient and flexible decoder that natively supports any arbitrary experimental design and can be easily integrated into automated pipelines. As sequence applications diversify, decoupling barcode classification from downstream analysis tasks becomes increasingly desirable, both to simplify workflows and to encourage the development of community standards for benchmarking and best practices.

Sequencing platforms generally tie in demultiplexing as a preprocessing step that is not directly accessible to users. The most widely used first-generation tool for sample demultiplexing is Illumina’s bcl2fastq, which combines basecalling with sample demultiplexing. bcl2fastq accepts input only in the proprietary Illumina BCL format and is specifically geared toward sequence data from Illumina platforms. It relies on a naïve decoding method based on exact string matching with up to one mismatch, in other words a simple minimum Hamming distance decoder, and quality assessment procedures that are not obviously accessible to end-users. bcl2fastq can also generate FASTQ files without demultiplexing, but most other tools that classify sequences using barcodes, such as Picard [9], also use simple minimum distance decoding.

Standalone, peer-reviewed, classifiers include deML [10], Bayexer [11], and Axe [12]. deML is capable of handling a single barcode set with either one or two segments. While it consults basecalling qualities and reports classification scores in Sequence Alignment/Map (SAM) [13] auxiliary tags, deML does not accurately reflect the probability of correct classification for two reasons: it estimates maximum likelihood only for the top candidates that are within a short Hamming distance, and it operates under restrictive assumptions that do not hold in most complex cellular indexing designs – i.e. that barcodes are uniformly distributed and that contaminating sequences are extremely rare. Bayexer attempts to train a naïve Bayes classifier by studying the error pattern when the insert sequence is shorter than the read length and thus provides a second observation of the barcode. Although this approach can potentially increase accuracy in those specific cases, it is not applicable to general barcode decoding and the tool fails to produce output when such redundancy is not present. Axe uses a set of pre-computed prefix trees to find a match within a given Hamming distance and can partially handle barcodes that differ in length. Although this method can be very fast, it ignores quality scores and is highly sensitive to upstream errors in the prefix, and so sacrifices accuracy.

None of the above tools account for the prior distribution of barcoded samples (which is often nonuniform) or explicitly account for noise (i.e. contaminating or indeterminate sequences), nor do they compute or report the posterior classification probability for individual reads. They also lack the ability to address arbitrary offsets within read segments, decode barcodes with more than two components, or handle multiple types or combinations of barcodes, and thus often require custom pre- or post-processing.

Pheniqs overcomes many of the limitations of other decoders by taking into account key aspects of current decoding requirements: scalability, assumptions about the number of tags and their location, integration of richer metadata, and the ability to explicitly account for basecalling quality scores, uneven barcode distributions, and presence of indeterminate sequences.

## Implementation

Pheniqs (**PH**ilology **EN**coder w**I**th **Q**uality **S**tatistics, pronounced *phoenix*) combines a generic and extensible approach to barcode decoding with flexible configuration options that easily accommodate custom experimental designs. It implements both a standard minimum distance decoder (MDD) based on Hamming distance and a probabilistic decoder (PAMLD). PAMLD consults basecalling quality scores as well as priors to compute the full posterior decoding probability for classification, rather than a simple maximum likelihood estimate.

### Decoding with the posterior probability

Classification based on barcodes involves extracting a subsequence *r* from an observed read, along with the basecall quality scores associated with the individual nucleotides in *r*, and decoding the original sequence *s*. Let *r* ∈ {*A,C,G,T, N*}^*n*^ be an observed sequence of length *n* extracted from the read and 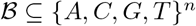 a given set of distinct barcodes where each 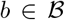 identifies an individual class. A decoder is denoted as a decision function 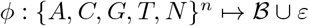, where *ε* is a decoding failure for an indeterminate sequence 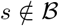.

A maximum likelihood decoder will identify the barcode 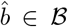 which maximizes the posterior probability that 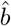 was sequenced given that *r* was observed.

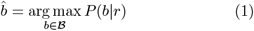

Applying Bayes’ rule we can compute *P*(*b*|*r*) using

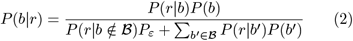

where *P_ε_* is the prior probability of encountering indeterminate sequences and 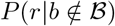 is the probability of observing a particular such sequence.

The *Phred-adjusted maximum likelihood decoder* (PAMLD) implemented by Pheniqs solves **Equation 2** by computing *P*(*r*|*b*) for each 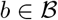 from the basecalling quality scores [14]. *P*(*b*), the expected fraction of reads identified by *b*, can be either provided *a priori* by the user or estimated directly from the data. In the absence of any prior information about potential sequence composition (base distribution or GC bias), we can only assume indeterminate sequences occur with maximum entropy so 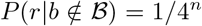. Such reads may arise from spiked in controls used for instrument calibration, contamination during library preparation or other unknown factors such as defective sequencing kits. Realistically, however, not every sequence in {*A, C, G, T*}^*n*^ is chemically stable enough to appear in sequencing, indeterminate entropy is lower, and 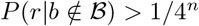. Empirical studies can determine a more refined lower bound for 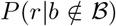. Pheniqs accommodates such refinements to the noise model by allowing advanced users to manually set this value.

When 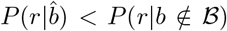, the initial evidence supporting the classification provided by the conditional probability is inferior to that provided by a random sequence, indicating that the 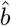 recovered in **Equation 1** cannot be distinguished from noise. The *noise filter* considers those a decoding failure without further consideration.

Reads that pass the *noise filter* are evaluated by the *confidence filter*, which compares 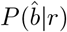 to a user-provided confidence threshold *C* for the minimum acceptable probability of a correct decoding and declares a failure if 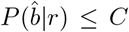. The probability of a decoding error is

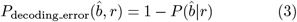

Directly estimating 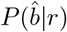 allows Pheniqs to report intuitive classification confidence scores for every read. Deriving a confidence score for a combinatorial barcode, made up of several independent components, requires to simply multiply the confidence scores of the individual components. The governing threshold C allows researchers to choose between assignment confidence and yield of classified reads and defaults to 0.95. The PAMLD decoding workflow is summarized in Figure 1B.

**Figure 1:**
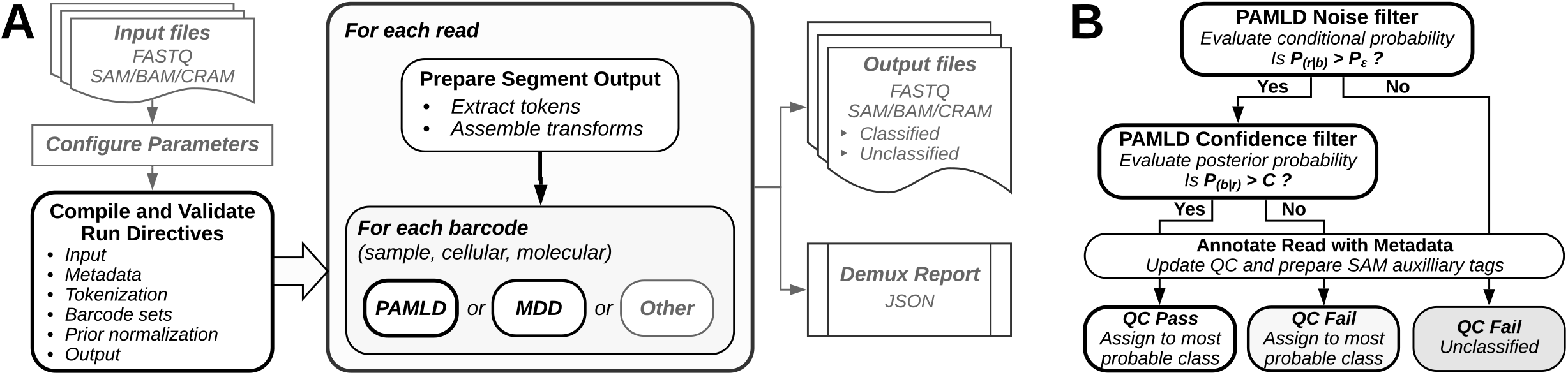
Pheniqs decoding pipeline architecture. ((A) Pheniqs requires as input sequence data files in any standard format and (if not using default parameters) a JSON configuration file. Python API tools (not shown) assist with IO management and can automatically generate an initial configuration using metadata from an Illumina run directory. Pheniqs evaluates each configuration component to determine how the data should be processed and to ensure that all required directives are present and properly specified. Any validation failures trigger clear, explicit error messages. Prior distributions of expected barcodes either derive from initial sample proportions as given (e.g. per a sample sheet), or are estimated directly from the data during a preliminary PAMLD decoding run. Barcode *tokens* are extracted from read segments using transform directives and then passed to a decoder (PAMLD or MDD). New decoding algorithms may be implemented as derived classes. Decoded barcodes and quality scores are written to specific SAM auxiliary field for each barcode type. Pheniqs can emit FASTQ files split by sample barcode, but SAM format is preferred since it preserves all associated metadata, and binary (BAM) and compressed (CRAM) versions produce considerably smaller files. POSIX integration allows direct piping to automated workflows, and support for real-time translation of file formats enables teeing to multiple outputs (thus avoiding the need to write temporary files). A JSON-encoded run report is also generated that provides summary statistics for the analysis. (B) PAMLD noise and confidence filters. Reads with a lower conditional probability than random sequences fail the *noise filter* and are classified as noise without further consideration. Reads with a posterior probability that does not meet the confidence threshold fail the *confidence filter*; these reads are classified, but they are marked as *qc fail* so the confidence threshold can be reconsidered at a later stage.

By contrast, deML [10] assumes that *P_ε_* is infinitesimally small and that samples are uniformly pooled (and thus equally likely), thereby suggesting that for every 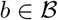

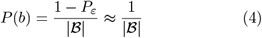

Under such conditions *P*(*b*|*r*) ∝ *P*(*r*|*b*) and **Equation 1** can be simplified to

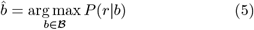

While such assumptions simplify maximum likelihood estimation of 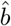, they are often grossly imprecise. For example, the relatively low yield in (e.g. single-cell) experiments that rely on several layers of combinatorial indexing often results in an extremely uneven barcode distribution, with *P_ε_* representing a significant portion of the sequenced DNA. Furthermore, implementations that refrain from computing 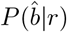 can not report the posterior classification probability or a confidence score for combinatorial barcodes.

### Estimating the prior distribution

Statistics from a preliminary PAMLD decoding run can be used to estimate the relative proportions of the individual barcodes *P*(*b*) for each 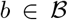 and the noise *P_ε_* from the data. The *high confidence estimator* bundled with Pheniqs estimates the relative proportions from the *high confidence* reads alone, assuming that *low confidence* reads (those that passed the *noise filter* but failed the *confidence filter*) and *high confidence* reads (those that passed both filters) come from the same distribution.

Let *S_ε_* be the number of reads rejected by the *noise filter*, *S_b_* the number of reads classified to *b* with confidence higher than *C*, and 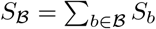. A *high confidence estimator* for the noise prior is

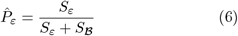

and for an individual barcode is

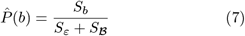

The *high confidence estimator* is a general purpose estimator, but Pheniqs can be used with any prior estimates devised by the user.

### Software Architecture

Pheniqs is distributed as a compiled binary with a command-line interface that accepts a JSON encoded configuration file (Figure 1A). Stable releases are also packaged in BioConda. To accommodate a rapidly changing landscape, Pheniqs is designed to be easily extensible with alternative error models and additional decoder implementations. An efficient codebase with robust input validation and comprehensive documentation supports individual installations and large-scale production facilities alike.

Using a familiar syntax that mimics Python array slicing, Pheniqs can decode multiple barcodes located anywhere in any sequence read. It extracts tokens from multiple read segments by addressing either the 5’ end, 3’ end, or both (and optionally reverse complemented) to construct the output template segments and the sample, cellular and molecular barcodes (Figure 2). For convenience, Pheniqs provides reusable definitions for several standard sample barcode sets that can be imported into any configuration file. Additional types of barcodes can easily be defined by declaring them in the configuration file (e.g. barcodes for split-pooling, antibody tags, spatial sequencing, etc.). This generic approach allows for arbitrary manipulation of sequence tokens that accommodates any potential barcoding scheme and obviates the need for pre- and post-processing for most experimental designs.

**Figure 2:**
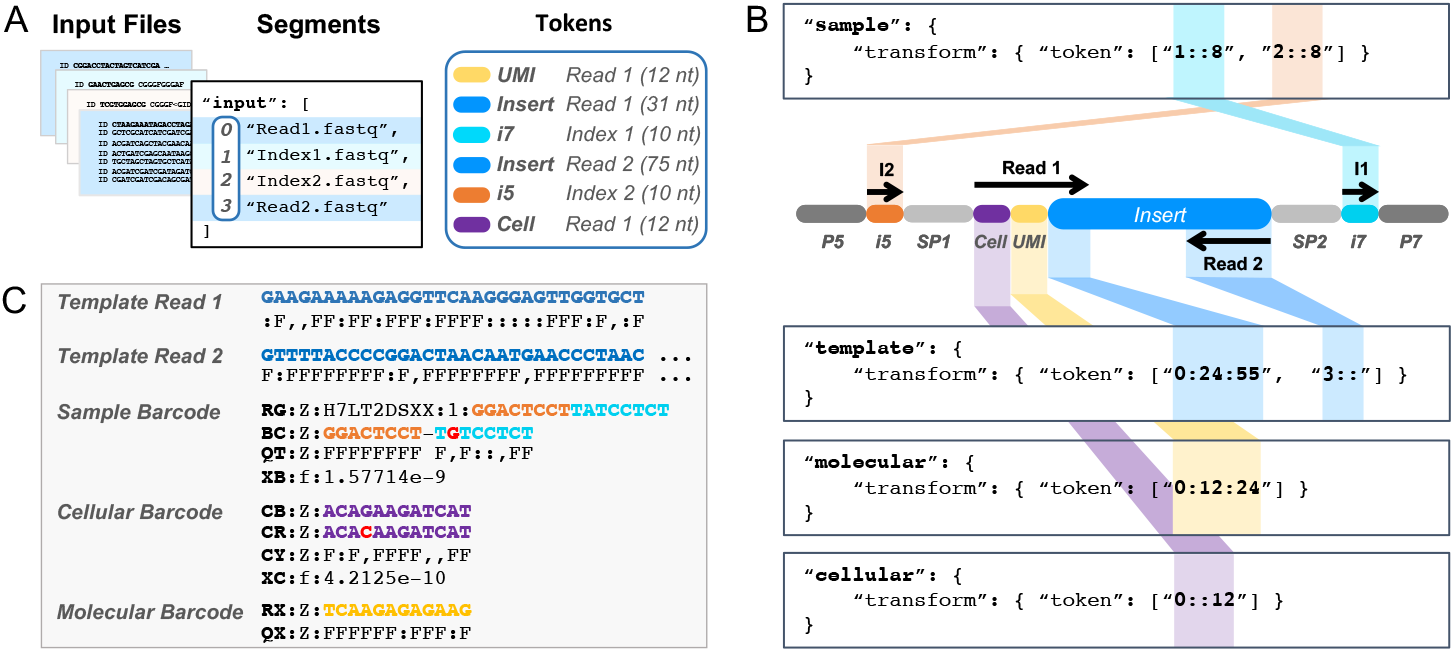
Tokenizing a standard Illumina read. Example of tokenization syntax for a 150-nt paired-end dual-indexed sequencing run. (A) Input files containing read *segments* emitted by the sequencer are indexed as an array, where 0=Read1, 1=Index1, 2=Index2, 3=Read2. Barcode *tokens* are defined for each type of barcode included in the experimental design and may appear at any position and orientation in any read segment. (B) This example contains a *sample* barcode composed of two 8nt elements (i5 and i7), a 12nt inline *cellular* barcode (Cell), and a 12nt inline *molecular* barcode (UMI). The biological sequences of interest (*template*) are located in Read 1 (31nt just downstream of the Cell and UMI) and all of Read 2 (here, 75nt). Each *token* comprises three colon separated components, “segment:start:end”. Per Python array slicing syntax, the *start* coordinate (offset) is inclusive and the *end* coordinate is exclusive. Start and end coordinates default to 0 and the end of the segment, respectively. (C) Template read segments, observed and most likely inferred barcode sequences, quality scores, and error probabilities are emitted to designated SAM fields. The reported error for each barcode is one minus its estimated confidence (the posterior decoding probability); for a combinatorial barcode, the reported error is one minus the product of the confidence scores for each component.

By directly interfacing with the low level HTSlib[15] C API Pheniqs can read, write, and manipulate either uncompressed or gzip compressed FASTQ files as well as the SAM file format or its binary compressed variants BAM and CRAM. Unlike FASTQ, the SAM format can encode sequencing data in a single, smaller file that supports richer metadata annotations. To allow multithreaded performance to scale linearly with core count, Pheniqs carefully synchronizes the many threads that read, decode and write with a consumer/producer model [16]. This allows threads that compute the posterior probabilities to work without waiting for threads that read and write. Each input feed uses two independent memory buffers: one for accepting incoming reads from the input file and one for supplying reads to the barcode decoding threads. When the first is full and the second is empty, a special thread momentarily locks both buffers and switches between them so that all operations may proceed with no interruption. The same principle is mirrored for output files. Since buffers may only be modified by one operation at a time to prevent data corruption, this design allows Pheniqs to concurrently receive input from multiple files, decode barcodes with multiple threads, and write output to multiple files, all with optimal efficiency. When integrated into a pipeline, Pheniqs can take advantage of POSIX standard streams to avoid the speed and storage bottlenecks associated with reading and writing temporary files.

Pheniqs reports the decoded sample, cellular, and molecular barcodes as well as their corresponding quality scores and the posterior decoding error probability in SAM auxiliary fields. It can associate standard SAM read groups with sample barcodes and can optionally perform an exhaustive quality assessment during processing that it includes in the final report.

## Results and Discussion

We used a variety of performance metrics to evaluate the decoding accuracy and computational efficiency of Phred-aware maximum likelihood decoding with Pheniqs (PAMLD), simple maximum likelihood estimation with deML, and minimum distance decoding (MDD, as implemented by Pheniqs). We used a semi-synthetic shortread dataset with a known true barcode set to measure accuracy across a range of error rates, and data from one lane of an Illumina NovaSeq run (containing >11 billion reads) to compare run time and memory usage.

### Accuracy

Decoding accuracy was analyzed using semi-synthetic barcoded sequence reads generated from the run published with deML [10]. To establish a ground truth for testing purposes, we simulated barcoded sequence reads by replacing the barcode nucleotides in each read with a perfect barcode sequence sampled from a known prior distribution. To simulate noise, we replaced a barcode with a sequence from a random offset in the genome of *PhiX174*, a 5386nt DNA bacteriophage with 45% GC content [17]). We used *PhiX174* sequences because Illumina sequencers require balanced and random nucleotide composition for instrument calibration and quality control, and *PhiX174* DNA is often spiked in as a control for this purpose at concentrations of 1%-5% or up to 40% for low-complexity libraries. *PhiX174* reads do not carry barcodes and should not be classified but instead labeled as Undetermined during demultiplexing. However, *PhiX174* sequences can sometimes contribute to noise contamination because they resemble an expected barcode by chance and must be removed in a downstream step by read sequence alignment. Notably, a recent study has found that 1000 genomes in the Integrated Microbial Genomes Database are contaminated with *PhiX174* sequences [18], suggesting that these are a common source of noise in Illumina sequence data.

We simulated sequencing errors by introducing a substitution at a nucleotide according to the basecall quality score and substitution frequencies made available with LRSim [19]. For this analysis we simulated substitution errors only since current short read platforms generate indel errors at a much lower rate than substitutions [20]. Finally, to simulate reads with different overall error rates, we recalibrated the quality scores produced in the first step and then simulated substitution errors on the recalibrated data. Figure S1 shows the calibrated quality score distributions of each simulated dataset. The modular architecture of Pheniqs allows for the addition of alternative error models that may be better suited for sequencing platforms with different error characteristics and could be tested similarly.

We used the above datasets to evaluate classification accuracy across a range of error rates for Pheniqs MDD with default settings (MDD); deML with default settings (deML); and Pheniqs PAMLD with default settings (PAMLD uniform), true priors (PAMLD true), and *high confidence* estimated priors (PAMLD estimated). Pheniqs can compute estimated priors in a preliminary run by updating an initial set of priors using observed barcode frequencies (Figure 3A shows the true sample barcode distribution in the dataset). *High confidence* estimated priors were computed here using statistics from a preliminary PAMLD run with the default 0.95 confidence threshold, 0.05 noise, and uniform barcode priors as input. We evaluated each barcode and the noise class as a binary classifier, so a correct assignment was counted as a true positive (*TP*), while an incorrect assignment was counted as a false negative (*FN*) for the correct class and as a false positive (FP) for the incorrectly assigned class. Reads marked as failing quality control (Figure 1B) were classified to the noise class for this analysis. We then summed up the values from all classes and computed the *false discovery rate* (FDR), *miss rate* (MR) and *F-score* (harmonic mean of the precision and recall).

**Figure 3:**
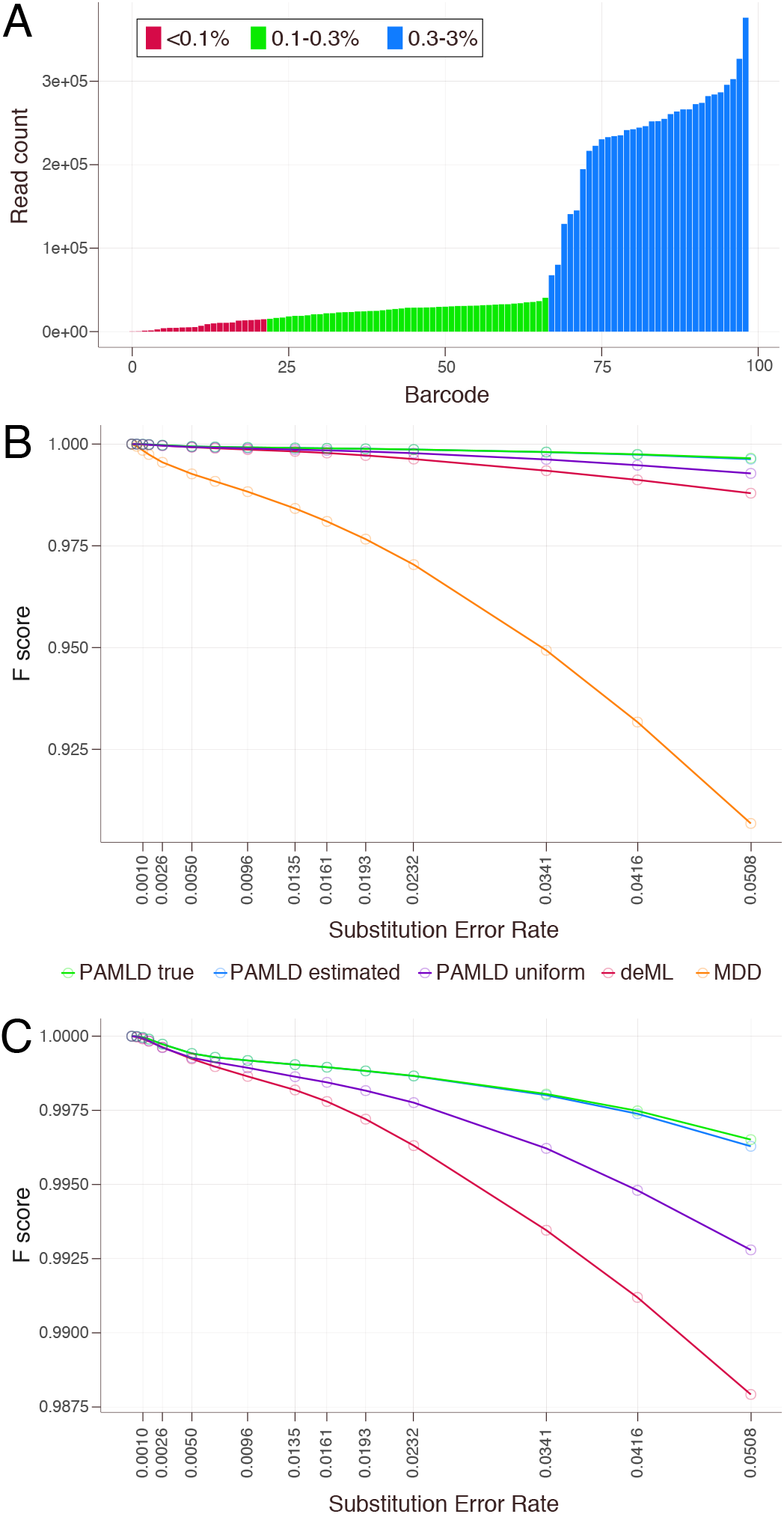
Decoding *F-score*. (A) True frequency distribution of individual barcodes in the synthetic data. The *control* barcode (the only one >3%, comprising 32% of the data used for evaluation) and *noise* (9%) were omitted for visual clarity. (B) All probabilistic decoders outperform minimum distance decoding by a wide margin, with differences becoming more pronounced as the error rate increases. (C) Among the probabilistic decoders, all configurations of PAMLD are more accurate than deML. The *high confidence* estimated prior performs nearly as well as the true prior.

The resulting analysis showed that the probabilistic decoders consistently outperform MDD, which has very little resilience to errors and noise (Figure 3B). Differences in accuracy become more pronounced as the substitution rate increases: at a rate of 0.05, the F-score for MDD is nearly 10% lower than other decoders. PAMLD with uniform priors is more accurate than deML, and *high confidence* prior estimation provides further gains that closely approximate performance with the true prior (Figure 3C). Thus, computing the full posterior probability outperforms simple maximum likelihood estimation, even under the same assumption of a uniform prior, and estimating the true prior distribution approaches near-optimal decoding.

The effect of the *noise filter* is illustrated in Figure 4, which shows the error in *high confidence* prior estimation for true barcode classes binned by their relative abundance across a range of substitution error rates. The difference between true and estimated priors is negligible at low overall error rates, but as basecalling quality decreases it becomes difficult to distinguish true barcodes from noise since *P*(*r*|*b*) for a true barcode is more likely to fail the *noise filter*. As a result, individual barcode classes are increasingly underestimated and 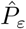 is correspondingly overestimated.

**Figure 4:**
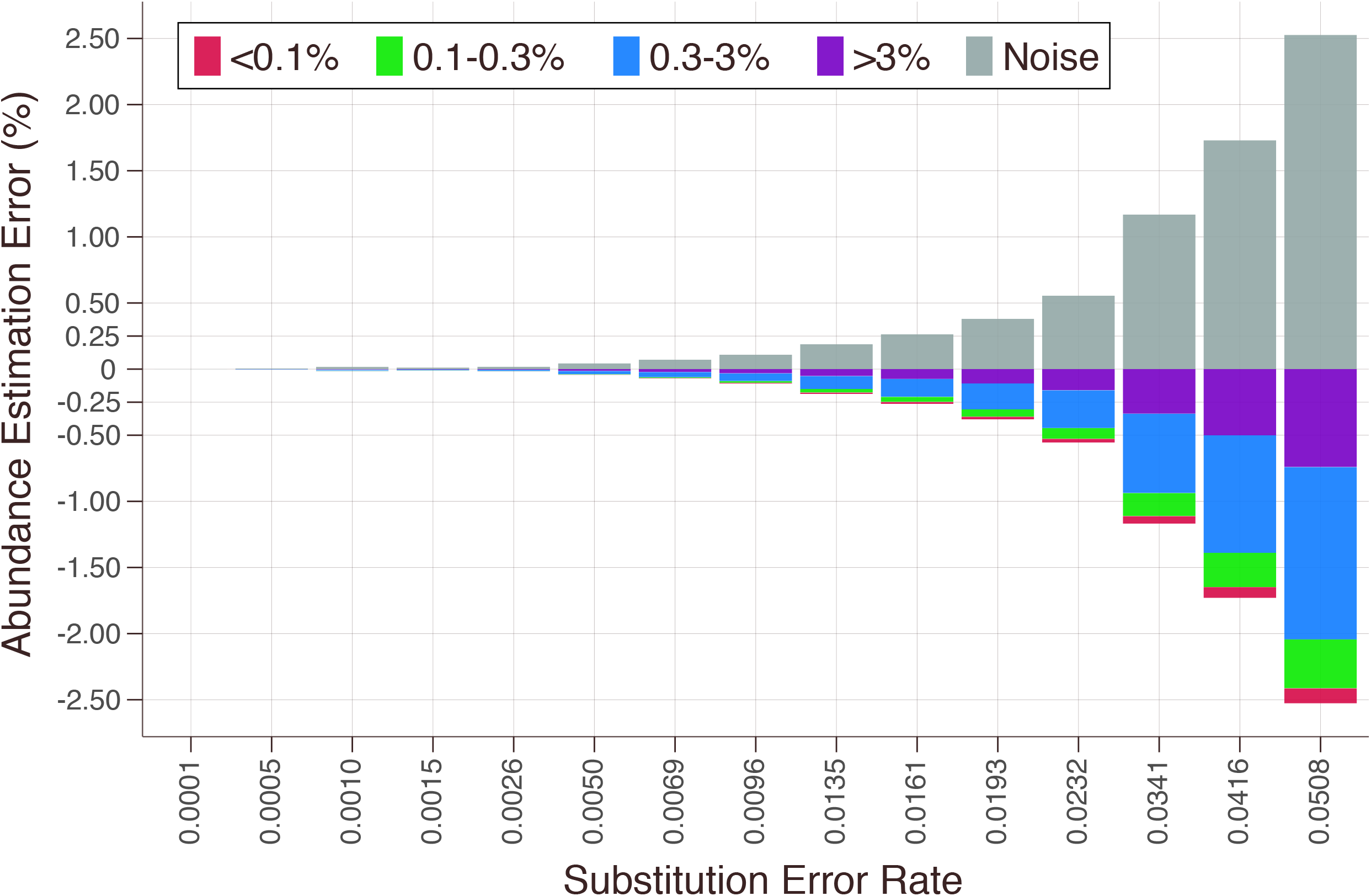
Error in high confidence prior estimation for true barcodes at different overall error rates. Results binned by relative barcode abundance reveals the tradeoff between noise and correct classification. As basecalling quality deteriorates, distinguishing true barcodes from noise becomes increasingly difficult, resulting in underestimation of the true barcode priors and overestimation of the noise prior 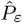.

To examine the sources of performance gains at a more fine-grained level, we plotted FDR, MR and *F-score* statistics separately for *classified*, *classifiable*, and *unclassified* reads across a range of error rates. *Classified* reads represent true barcodes or noise reads that were classified to a real barcode. These include correct assignments (TP), noise or barcode reads assigned to the wrong barcode class (FP), and true barcodes not assigned to their proper class (FN). While all decoders perform reasonably well on *classified* reads overall (Figure S2), Pheniqs with true or estimated priors consistently results in a lower rate of misclassified reads (lower FDR) and recovers more reads (lower MR) than other decoders. Filtering noise helps Pheniqs outperform deML in any configuration, primarily by reducing FDR (Figure S2A). In the lower error range relevant to most Illumina runs (~1/1000), PAMLD shows a lower FDR than even MDD (Figure S2B). The FDR is flat for MDD because it ignores any reads with more than one mismatch, and for the same reason the miss rate climbs dramatically relative to the other decoders as the error rate increases.

Since datasets now contain increasingly large numbers of multiplexed samples or individual cells, the ability to accurately identify rare barcode classes is of high interest. We used read counts for each barcode class (Figure 3A) to examine performance for *classified* reads binned by their relative abundance: *very low* (<0.1% of reads), *low* (0.1% to 0.3%), *similar to uniform* (0.3% to 3%) and *overrepresented* (>3%; here a single control barcode accounted for 32% of all reads). Across the entire spectrum, PAMLD with estimated or true priors shows better overall performance (F-score) than either deML or PAMLD with uniform priors (Figure S3). For more abundant barcode classes, gains are mainly due to higher sensitivity (lower MR). For lower abundance barcodes (<0.3% of total reads), PAMLD makes ~10-fold fewer incorrect assignments than deML (lower FDR) with only a modest loss of sensitivity. Thus PAMLD is especially beneficial for the detection of rare barcodes.

Indeterminate barcodes are not uncommon in shortread datasets and can greatly reduce the yield of usable data. We found that PAMLD improves accuracy for both *classifiable* reads (true barcode reads, correctly classified or not) and *unclassified* reads (reads that are noise or fail quality control). For *classifiable* reads, both FDR and MR are consistently lower, leading to improved F-scores (Figure S4; examples in Figure 5A-C). For *unclassified* reads, probabilistic decoders have a lower FDR (Figure S5) and also classify many fewer true barcodes as noise than MDD (Figure S5; examples shown in Figure 5A,B,D). The PAMLD *noise filter* rejects reads when the evidence for the barcode with the highest posterior is no better that for a random sequence (Figure 1B). This is arguably a desired property since such reads have very weak evidence for classification. When the overall error rate is low, PAMLD errs on the side of caution and classifies slightly more true barcodes as noise than deML (FDR, Figure S5A). On the other hand, PAMLD misses many fewer true noise reads (lower MR) than deML (Figure S5A; example shown in Figure 5D), and in the range of error rates for Illumina sequencers filters true noise better than even MDD (lower MR, Figure S5B).

**Figure 5:**
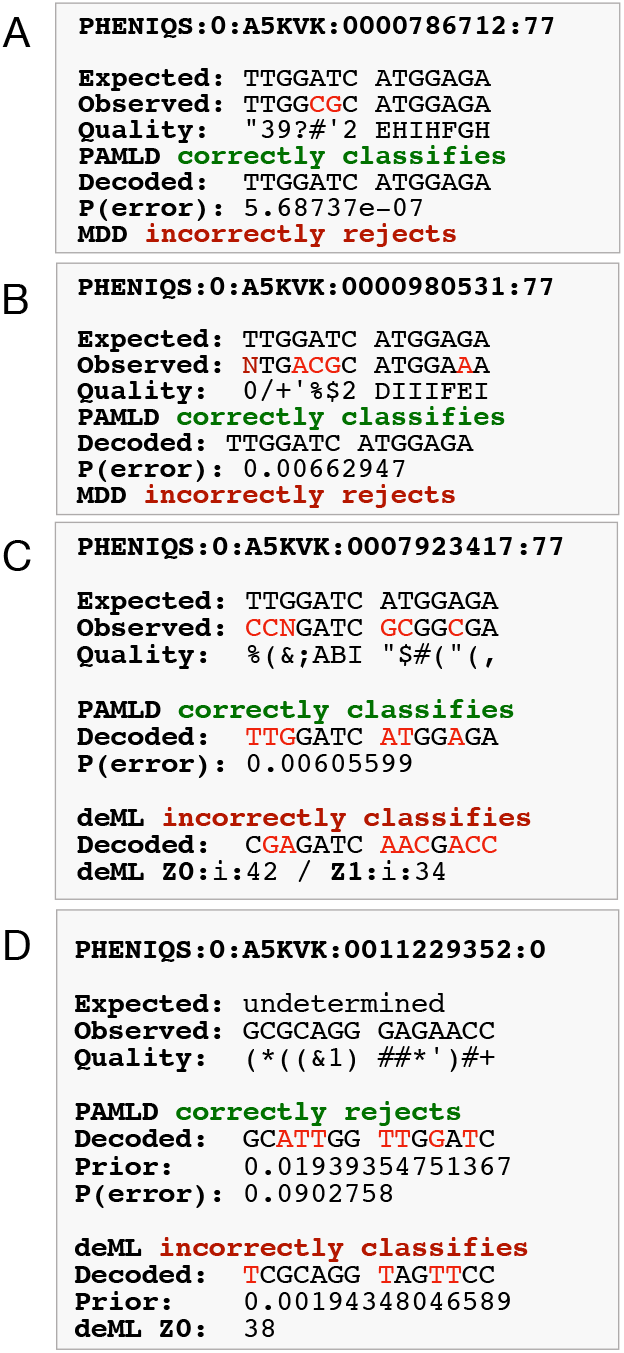
Decoding examples. (A) The barcode on this read has only two mismatches (highlighted in red in *Observed*). MDD rejects it and assigns it to the undetermined bin. PAMLD easily classifies it with a confidence of more than 99.9999%. This is a good example of why MDD is overly conservative and has such a high rate of false negatives. (B) This example has 4 mismatches and one uncalled nucleotide (highlighted in red in *Observed*). MDD rejects it while PAMLD still classifies it with relatively good confidence of 99.337%. (C) In this example there are 5 mismatches and one uncalled nucleotide (highlighted in red in *Observed*). PAMLD classifies it correctly with a good confidence of 99.3944%, while deML classifies the read to the wrong bin. Mismatched nucleotides in the *Observed* sequence are marked in red. (D) In this example the read was noise and should have been rejected and assigned to the undetermined bin. PAMLD initially classifies it to a barcode with 6 mismatches, but it rejects the classification because the error probability is above 9%. deML incorrectly classifies the read. PAMLD’s decision is influenced by the prior for the barcode it chooses, which is about 10 times higher than the barcode chosen by deML.

In summary, we find that using prior knowledge of barcode frequency and sequence quality statistics, combined with noise filtering, increases the recovery of true barcodes and reduces the number of misclassified reads. These benefits are most pronounced for barcodes present at low frequency and for data with high error rates. Thus probabilistic, quality-aware decoding offers distinct advantages for classifying short-read datasets that are highly complex and/or use combinatorial barcoding strategies. Pheniqs may also prove advantageous for third-generation single-molecule sequencing platforms that provide long sequence reads at much higher error rates.

### Runtime Speed and Memory

To evaluate speed and memory usage we used a single lane from an Illumina NovaSeq, the highest throughput instrument available today, containing 94 multiplexed libraries with standard dual i7 and i5 barcodes. 151 cycles were sequenced on each of the two template segments and 8 cycles on each of the i7 and i5 index segments.

Benchmarking for speed (Figure 6A) and memory (Figure 6B) was executed on an Intel Xeon CPU E5-2690 v4 @ 2.60GHz with 14 cores and 28 threads. Basecalling the lane with bcl2fastq (without demultiplexing) took 47 minutes and yielded 11,578,868,372 dual-indexed paired-end reads that passed quality filtering. bcl2fastq produces reads with segments split over four gzip compressed FASTQ files that were used as input to Pheniqs. Pheniqs runtime benchmarks show that output format encoding greatly impacts speed. With null output Pheniqs decodes the barcodes and collects statistics (for instance to be used for prior estimation) but does not write any output. The null benchmark completed in less than half the time of all others, demonstrating that writing output files is the rate-limiting factor for performance. Pheniqs performs fastest when producing interleaved BAM output and demultiplexed the NovaSeq run in 3 hours and 10 minutes. Even computing the full posterior probability, Pheniqs scales almost linearly with the number of available cores and is at least 20 times faster than deML, which lacks support for multithreading. deML could only produce FASTQ output with FASTQ input, so we could not test BAM output.

**Figure 6:**
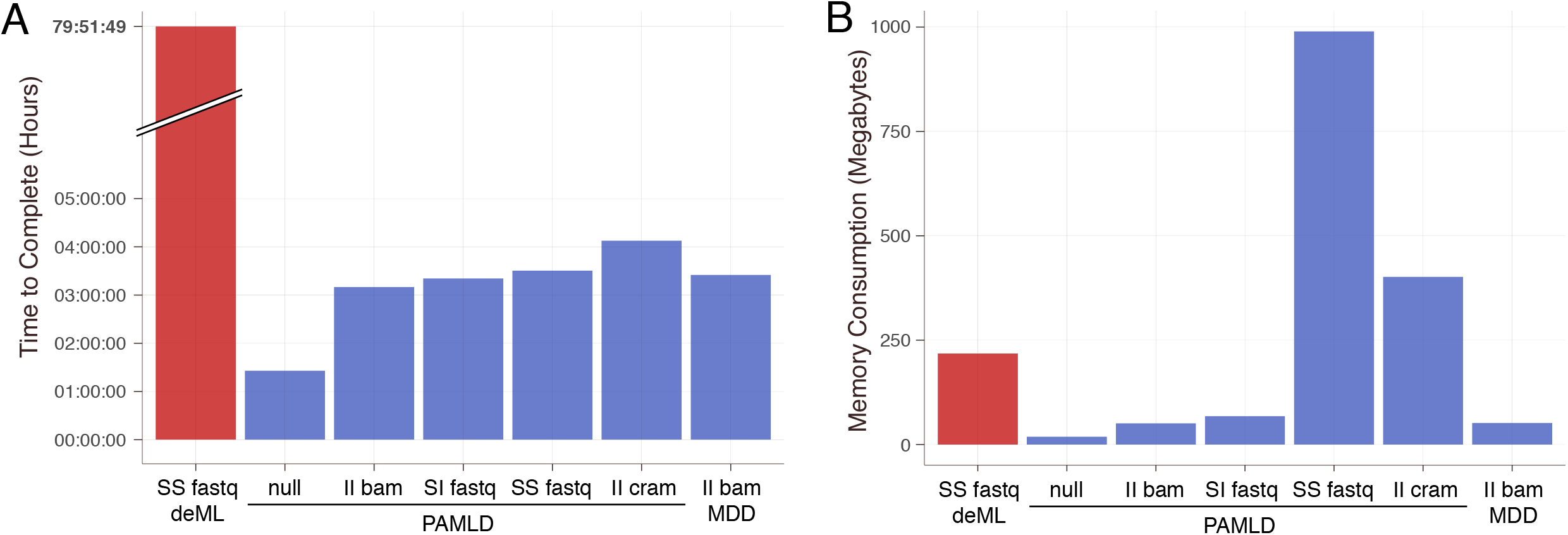
Speed and memory benchmarks. One lane of an Illumina NovaSeq run was analyzed. The two letter codes describe the output layout: S for *split* and I for *interleaved*, first for library and second for segment. (A) Pheniqs with PAMLD generated interleaved BAM output in 3:10 hours. CRAM is much more complex than BAM and employs several compression algorithms that are more aggressive than gzip to produce smaller files, so writing CRAM output took slightly longer as expected. Pheniqs scales almost linearly with the number of available cores and is at least 20 times faster than deML, which took almost 80 hours to complete. (B) Pheniqs memory consumption increases linearly with the number of output files, since each is allocated separate IO buffers, but even with 200 output files and generous buffers Pheniqs still used less than 1GB of memory.

## Future Work

We can envision a variety of ways to enhance the performance of Pheniqs. Our initial focus will be to facilitate integration into standard analysis workflows. To assist with configuration, we are building a library of barcode configuration templates for common sequencing kits and experimental designs. The Pheniqs website currently includes vignettes for configuring standard Illumina sequencing runs, single-cell RNA sequencing using the Fluidigm platform, and two plate-based split-pooling methods for single-cell RNA-seq: sciRNA-seq [2] and SPLiT-seq [3]. We also plan to add support for other types of single-cell and multimodal profiling, as well as novel sequencing applications as they arise.

One limitation of Bayesian decoding is that computing the posterior probability requires knowing the list of expected barcodes in advance. To overcome this problem, the same strategy used by Pheniqs to estimate priors for a known set of barcodes could be extended to an unknown set of barcodes. First, a preliminary run configured with a whitelist (e.g. a list of all known cellular barcodes in a single-cell sequencing kit) can be used to estimate the relative abundances of observed barcodes. A shorter list of barcodes can then be extracted from the preliminary run report by placing a threshold on the abundance of observed barcodes, which may then be decoded in a second run using the imputed priors. Providing decoding quality scores for whitelisted cellular barcodes based on the full posterior probability will allow improved estimation of their relative abundance, thus increasing accuracy and sensitivity for single-cell applications. An overall classification quality score combining multiple types of barcodes can then be computed, which will be easy to both report and understand. Since the complexity of computing the posterior is linearly correlated with the size of the barcode set, however, computing priors for a very large whitelist may become impractically slow. To address this, we are considering alternative strategies to reduce the complexity of the problem in such cases.

Another issue that Pheniqs currently does not address is error correction for UMIs (unique molecular identifiers) — barcodes that tag individual molecules in a library, which are used to identify PCR duplicates and to quantify the abundance of distinct molecular species. Since reads classified to a single combination of sample, cellular and molecular barcode are assumed to be a clone of the same biological sequence, the barcodes are not independent, making computing the full posterior probability unfeasible. To incorporate error correction for UMIs, we are looking into heuristic algorithms that rely on established peer-reviewed methods.

We are also interested in applying Pheniqs to data from other sequencing platforms. Performance evaluations of PAMLD on synthetic data show that gains in decoding accuracy become more pronounced as the rate of substitution errors increases. This suggests that Pheniqs could greatly enhance the analysis of sequence data with high error rates, such as third generation single-molecule sequencing platforms, which currently operate with error rates in the range of several percent [21] [22] [23]. In order to apply PAMLD to PacBio and Oxford Nanopore sequencing, we will need to incorporate support for insertions/deletion errors, which are more common on these platforms (particularly at homopolymers). We thus plan to evaluate performance and to investigate alternative noise models for these sequencing technologies. Finally, working toward common benchmarks would be useful for the development of community standards for barcode decoding quality, an issue that can have profound impacts on data quality but has not so far received widespread attention.

## Conclusion

We show that barcode classification using full posterior probabilities with noise filtering is more accurate than other available methods. We provide a multithreaded software package with comprehensive input validation, Pheniqs, that is faster and more scalable than existing tools. The probability model implemented by Pheniqs accounts for both erroneous codewords and non-codeword random noise, can handle arbitrarily complex barcoding designs, and generalizes to multiple combinatorial tags. It relies on intuitive confidence thresholds for fine-tuning decoding accuracy and reports decoding confidence scores and barcode sequences for individual reads by populating standardized SAM format auxiliary fields. Pheniqs is designed for integration into automated analysis workflows and can be extended with new error models and alternative decoders. An efficient implementation, coupled with comprehensive documentation and a suite of helper tools, supports both individual users and core facilities alike.

## Availability and requirements

- **Project name:** Pheniqs
- **Home page:** http://biosails.github.io/pheniqs
- **Bioconda:** https://anaconda.org/bioconda/pheniqs
- **Operating systems:** Linux, macOS.
- **Programming language:** C++11, Python 3.
- **Other requirements:** clang, gcc 4.8 or newer.
- **License:** NYU, free for academic use.
- **Restrictions to use by non-academics:** license needed.

## List of abbreviations

SAM: Sequence Alignment/Map format
MDD: Minimum Distance Decoding
PAMLD: Phred-adjusted maximum likelihood decoding
UMI: Unique Molecular Identifier. Also referred to a Molecular Barcode.
TP: True positive.
FP: False positive.
FN: False negative.
FDR: False discovery rate 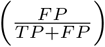.
MR: Miss rate 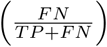.
F-score: harmonic mean of precision and recall.
nt: Nucleotide.
IO: Input / Output.

## Availability of data and materials

Synthetic data was generated using the method described in the text with data made publicly available with deML [10]. BAM file is available at https://bioinf.eva.mpg.de/deml

## Authors’ contributions

LG conceived the project, developed the algorithms, wrote the code, and co-authored the manuscript. DS inspired the probabilistic likelihood model, reviewed the mathematical formulation, and provided feedback on the manuscript. KCG supervised the project, motivated the methodology for combinatorial barcode decoding, and co-authored the manuscript with LG.

## Acknowledgements

We are grateful to Kapil Thadani and Or Biran for discussions on the statistical evaluation and mathematical notation; to Alan Twaddle for his feedback on the user manual and his help with benchmark analysis and figure preparation; to Jillian Rowe for her assistance with Conda packaging, and continuous integration; to Nizar Drou for constructive input and assistance with benchmarking; to Mohammed Khalfan for feedback on Pheniqs version 1 and for helpful discussions on the documentation; and to Giuseppe Saldi for his insightful comments.

