## supplemental materials for "Pheniqs 2.0: accurate, high performance Bayesian decoding and confidence estimation for combinatorial barcode indexing"

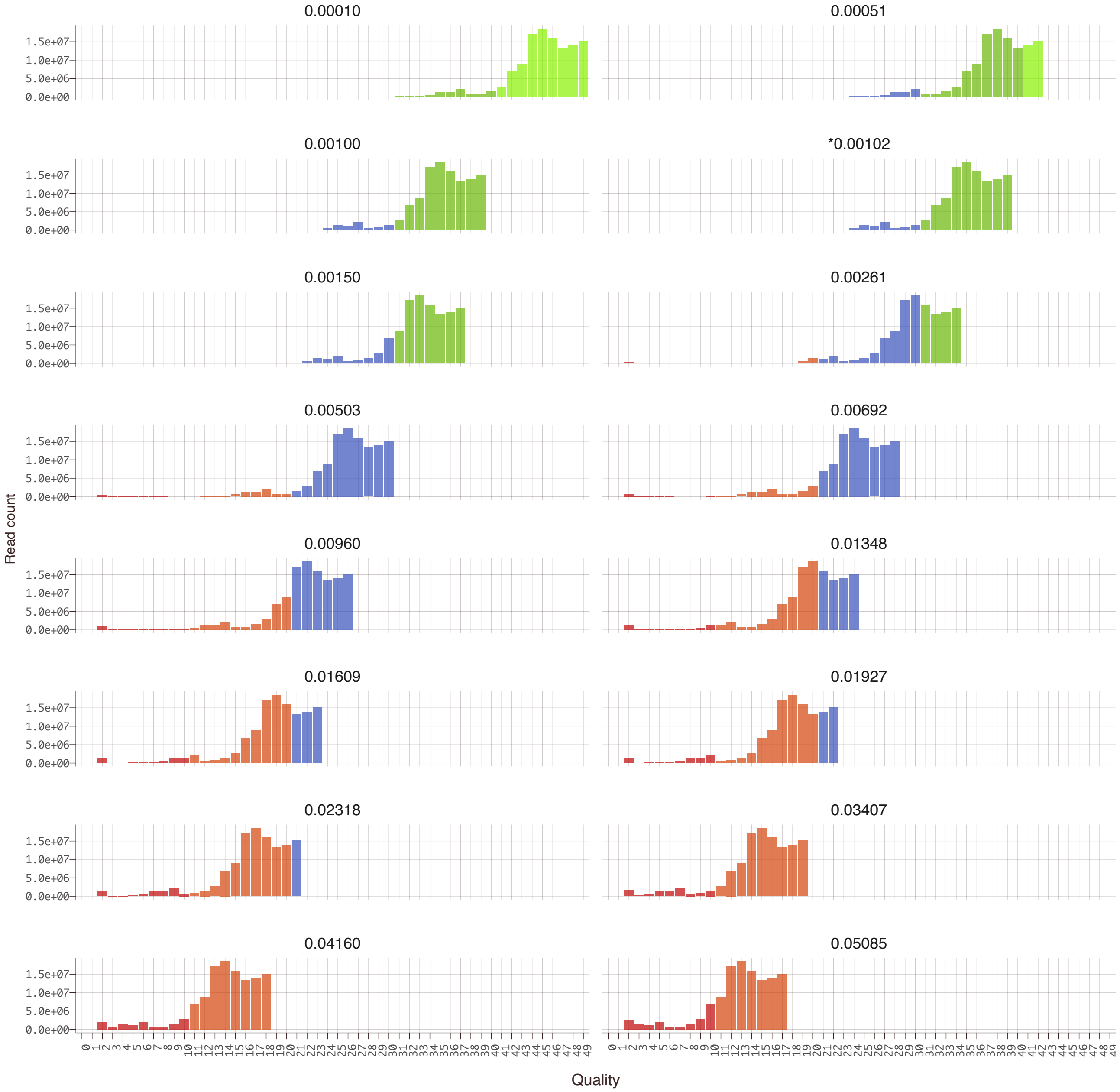


Figure 1: **Basecall quality distribution for barcode cycles.** Quality scores were recalibrated to yield different overall error rates, while preserving the histogram topology, by applying a linear factor. The different colors represent orders of magnitude on the Phred scale. The original uncalibrated distribution is indicated with an asterisk (overall error rate = 0.00102).

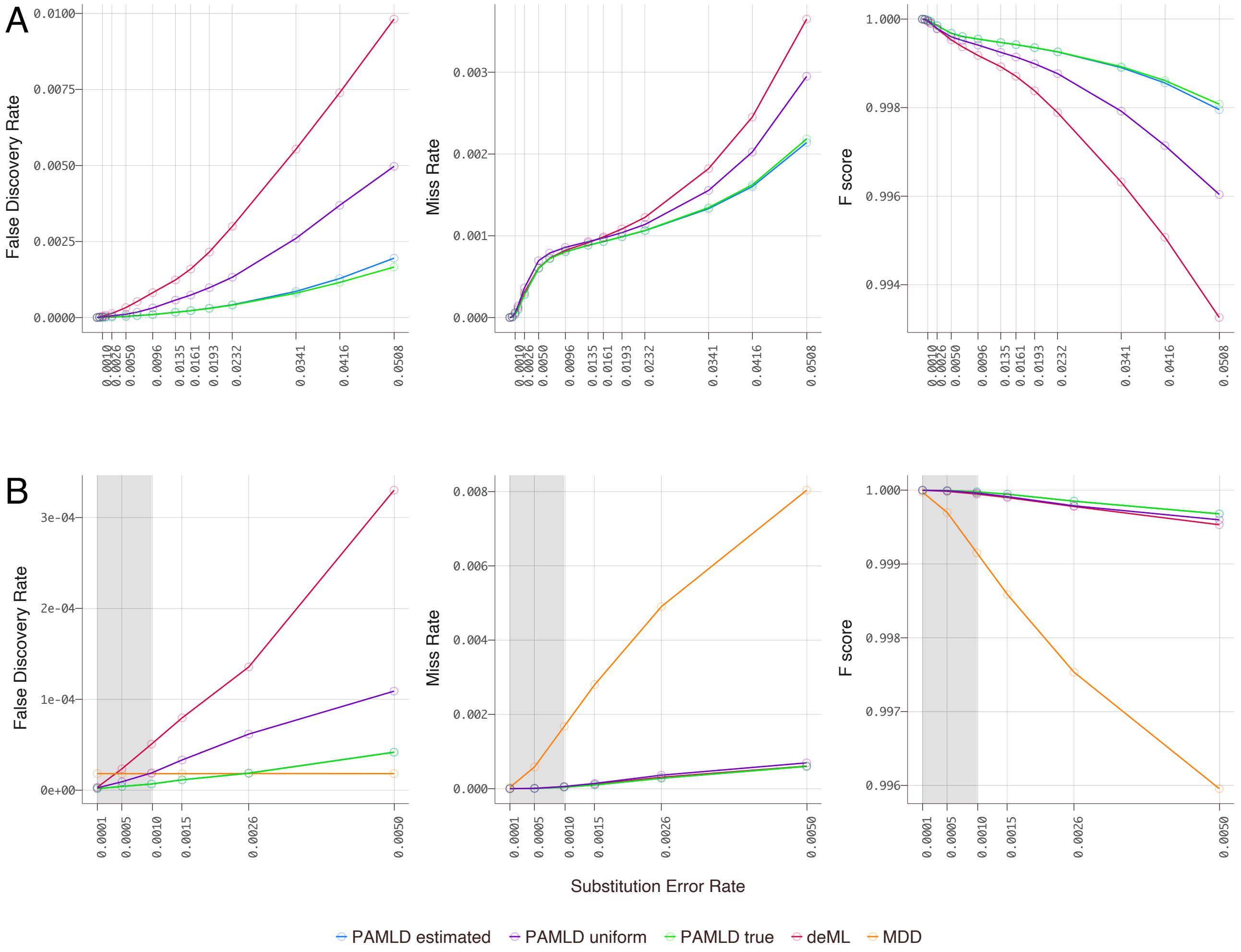


Figure 2: **Accuracy for *classified* reads only.** *Classified* reads are all reads (true barcodes or noise) that were classified to a barcode. (A) Pheniqs outperforms deML in every configuration. The improvement is more pronounced for FDR than for MR. (B) In the basecall error range relevant to most Illumina runs (on the order of $10^{-3}$), PAMLD has lower FDR than even MDD.

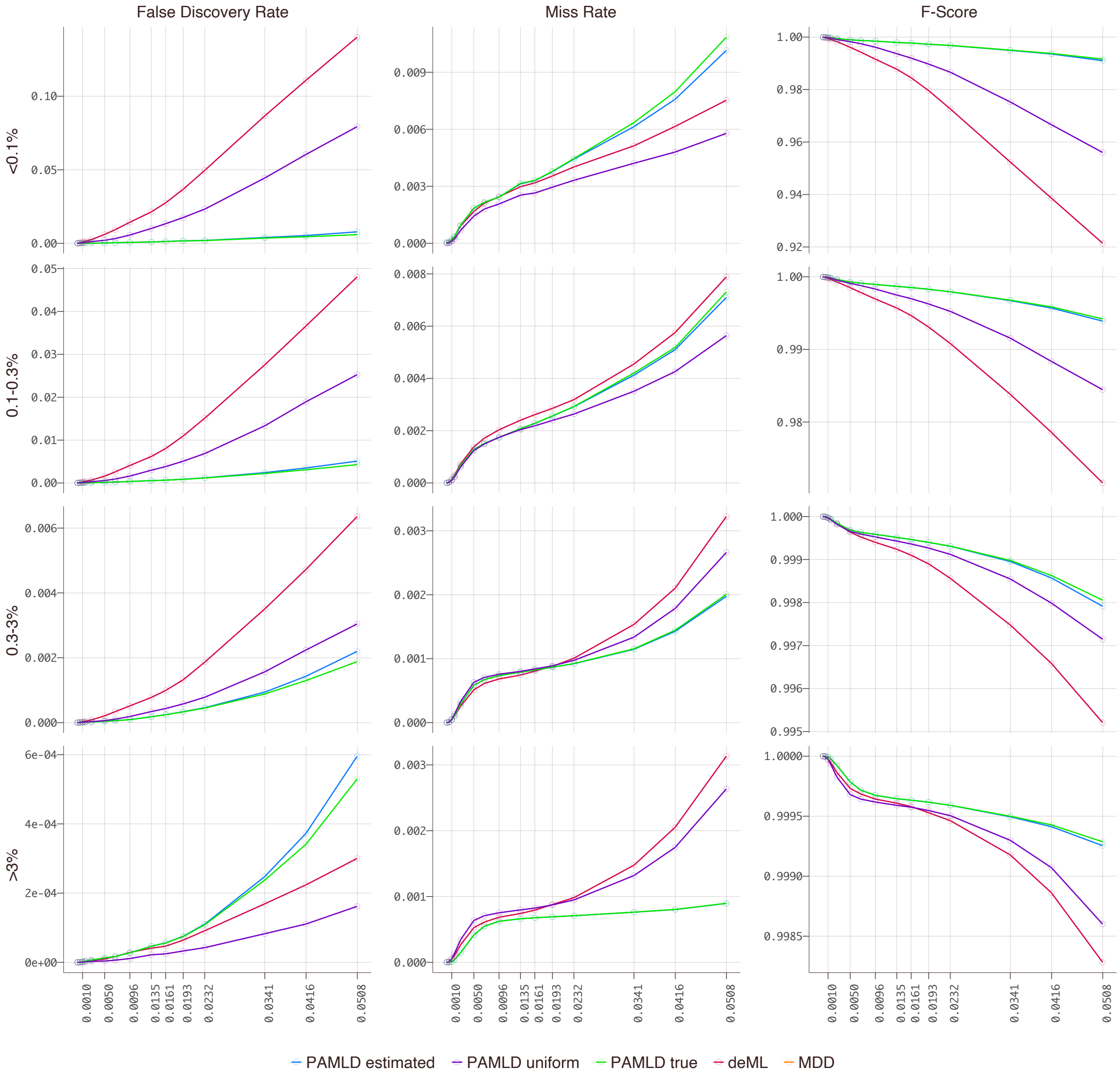


Figure 3: **Accuracy for classified reads by relative abundance of true barcodes.** Barcode classes were binned according to their true priors (as in Figure 5): $< 0.1\%$ (*very low abundance*), 0.1-0.3% (*low abundance*), 0.3-3% (*similar to uniform*), or $> 3\%$ of total reads (the *overrepresented* class contained only one barcode present at 32%). When classifying *very low abundance* reads, PAMLD with estimated or true priors reduces FDR to extremely low levels, with a minor tradeoff in MR. Reads in classes with a *low abundance* prior show a similar improvement in FDR and a slightly lower MR than deML. Prior estimation improves both FDR and MR for *similar to uniform* barcode classes. Reads from the *overrepresented* class show negligible (on the order of $e^{-4}$) FDR by any decoding method; notably, the MR for PAMLD with priors is about the same magnitude, whereas for deML the MR is around ten-fold higher than FDR. The F-score is consistently higher for PAMLD with estimated or true priors and becomes more pronounced with increasing error rate.

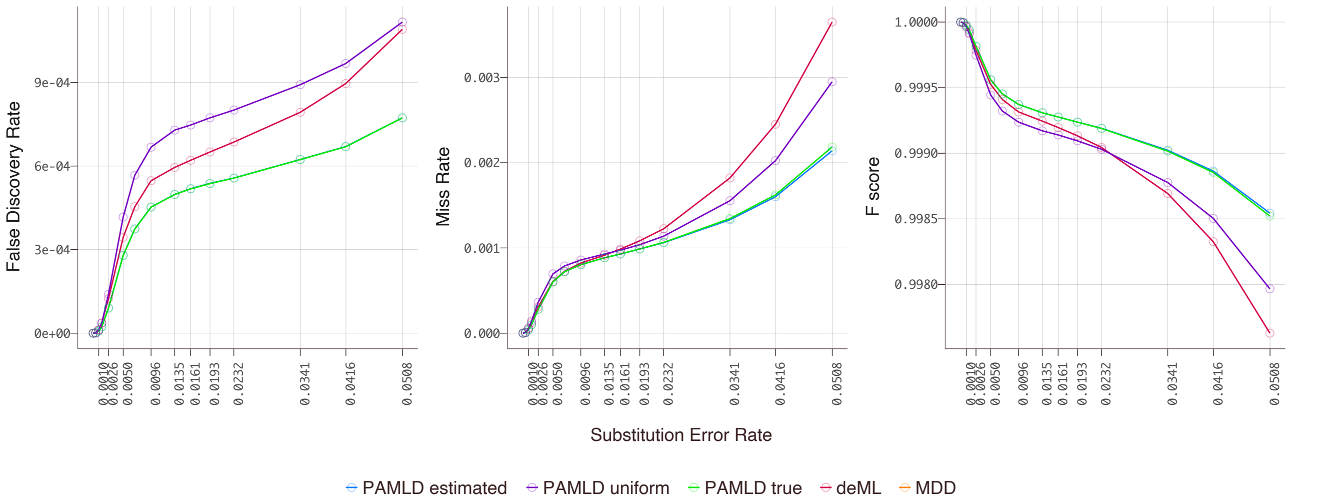


Figure 4: **Accuracy for classifiable reads only.** *Classifiable* reads are all true barcoded reads, correctly classified or not. Adding prior knowledge improves both FDR and MR, resulting in better overall performance.

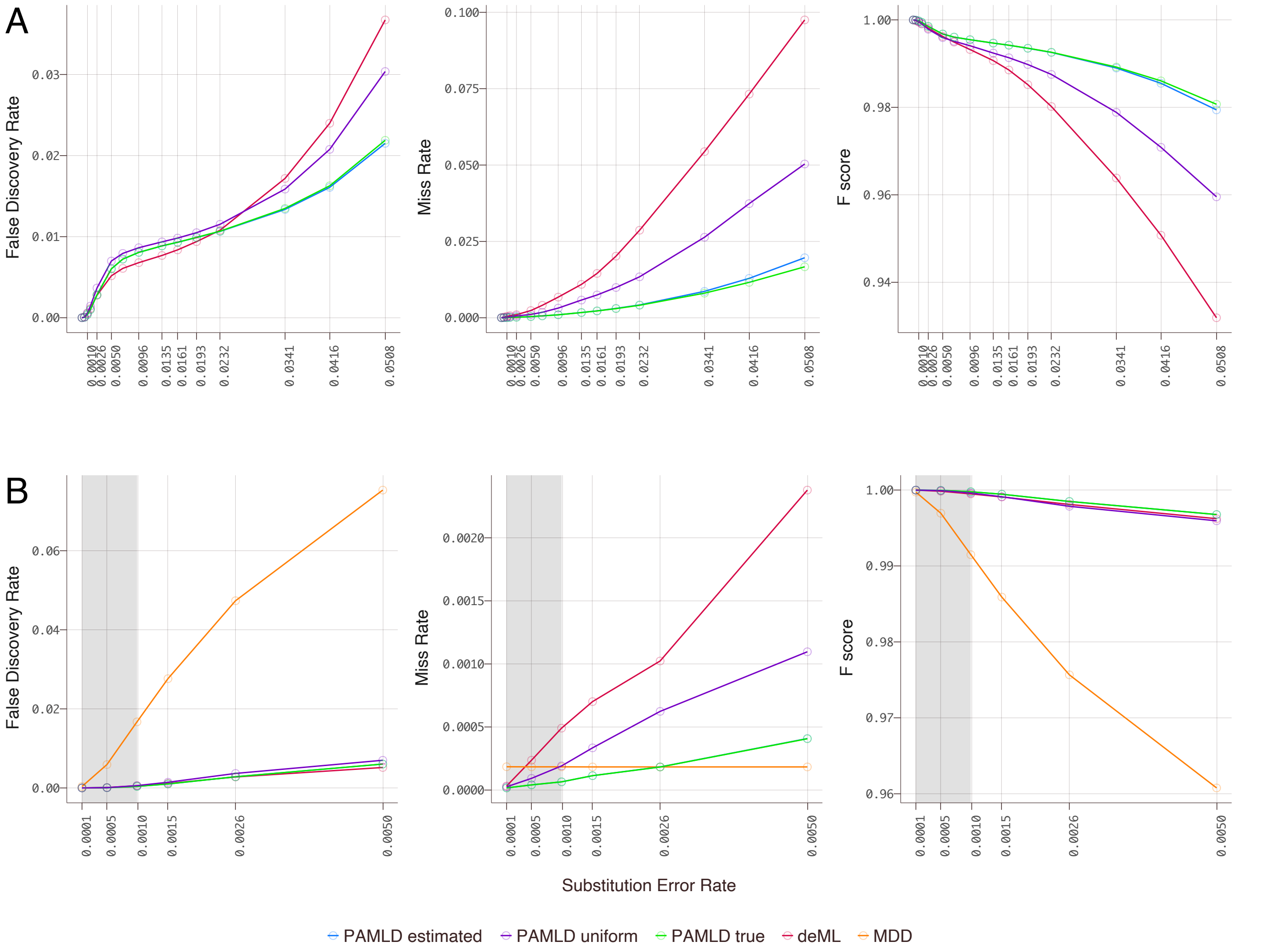


Figure 5: **Accuracy for unclassified reads.** (A) Comparison of probabilistic decoders. PAMLD decoders are more aggressive in detecting noise than deML and classify slightly more true barcoded reads as noise when the error rate is low. The *noise filter* will reject the read when evidence supporting the class with the highest posterior does not exceed that provided by a random sequence, even if the posterior is high. (B) Comparison with MDD for unclassified reads at low error rates. PAMLD misses very few noise reads, even outperforming the very conservative MDD in the range relevant for Illumina platforms.
